# High-throughput automatic training system for odor-based cognitive behaviors in head-fixed mice

**DOI:** 10.1101/255653

**Authors:** Zhe Han, Xiaoxing Zhang, Jia Zhu, Yulei Chen, Chengyu T. Li

## Abstract

Understanding neuronal mechanisms of cognitive behaviors requires efficient behavioral assays. We designed a high-throughput automatic training system (HATS) for olfactory behaviors in head-fixed mice. The hardware and software were constructed to enable automatic training with minimal human intervention. The integrated system was composed of customized 3D-printing supporting components, an odor-delivery unit with fast response, Arduino based hardware-controlling and data-acquisition unit. Furthermore, the customized software was designed to enable automatic training in all training phases, including lick-teaching, shaping, and learning. Using HATS, we trained mice to perform delayed non-match to sample (DNMS), delayed paired association (DPA), Go/No-go (GNG), and GNG reversal tasks. These tasks probed cognitive functions including sensory discrimination, working memory, decision making, and cognitive flexibility. Mice reached stable levels of performance within several days in the tasks. HATS enabled an experimenter to train eight mice simultaneously, therefore greatly enhanced the experimental efficiency. Combined with causal perturbation and activity recording techniques, HATS can greatly facilitate our understanding of the neural-circuitry mechanisms underlying cognitive behaviors.

## Introduction

Behavioral design and analysis are critical for understanding neural mechanism of cognition (Gomez-Marin et al., 2014), including working memory (Fuster, 1997; Baddeley, 2012), decision making (Gold and Shadlen, 2007; Lee et al., 2012), and reversal of learnt rules (Bunge and Wallis, 2008). Combined with novel neural-circuitry technologies, such as optogenetics (Fenno et al., 2011), chemogenetics (Armbruster et al., 2007), and imaging methods (Deisseroth and Schnitzer, 2013), well-disigned behavioral paradigms can greatly facilitate the ciruitry level understanding of behaivor. Reliable behavioral paradigms are also useful in pre-clinic studies such as target identification and mechanistic studies for brain diseases (Fernando and Robbins, 2011, Gotz and Ittner, 2008, Nestler and Hyman, 2010).

Optimally, behavioral training systems should be automatic, ready to scale up, blind in design, and flexible in changing paradigms. Automatic training systems (Schaefer and Claridge-Chang, 2012) met well with these criteria. There was a long history of designing automatic behavior-training systems, for example in studies of operant conditioning (*e.g.,* Davidson et al., 1971). Automatic training systems are composed of monitoring and feedback controlling of behavior. In free-moving mice, automatic measurement has been implemented in characterizing visual performance (Benkner et al., 2013, de Visser et al., 2005, Kretschmer et al., 2013), evaluation of pain sensitivity (Kazdoba et al., 2007, Roughan et al., 2009), freezing behavior during fear conditioning (Anagnostaras et al., 2010, Kopec et al., 2007), home-cage phenotyping (Balci et al., 2013, Hubener et al., 2012, Jhuang et al., 2010), anxiety (Aarts et al., 2015), diurnal rhythms (Adamah-Biassi et al., 2013), and social behavior (Hong et al., 2015, Ohayon et al., 2013, Weissbrod et al., 2013). With feedback controlling components, automatic training systems have been successfully implemented in multiple behavioral domains, including memory assessment (Reiss et al., 2014), operant learning (Remmelink et al., 2015), and training limb function (Becker et al., 2016). Automatic training systems with multiple cognitive behaviors requiring memory, attention, and decision making have been developed previously in free-moving rats (Erlich et al., 2011, Poddar et al., 2013) and mice (Gallistel et al., 2014, Romberg et al., 2013). Moreover, such systems were successful in dissecting neural-circuitry mechanisms underlying cognitive behaviors (*e.g.,* Brunton et al., 2013, Erlich et al., 2011, Hanks et al., 2015). Head-fixed mice (Dombeck et al., 2007; Guo et al., 2014) renders great flexibility in recording (Harvey et al., 2009; Boyd et al., 2012; Fukunaga et al., 2012; Kollo et al., 2014) and imaging (Boyd et al., 2015; Chu et al., 2016, Dombeck et al., 2007, Komiyama et al., 2010; Yamada et al., 2017) technologies. Moreover, free-moving and head-restrianted mice exhibit similar ability of olfactory discrimination (Abraham et al., 2012). However, automatic training systems in head-fixed mice were not developed previously.

Olfaction is an important sensory modality for cognitive behavior (Doty, 1986; Ache and Young, 2005). Previous studies have demonstrated that rodents are very good at olfactory discrimination, memory, and decision (Abraham et al., 2004, Barnes et al., 2008, Cleland et al., 2002, Haddad et al., 2013, Hubener and Laska, 2001, Kepecs et al., 2007, Komiyama et al., 2010; Liu et al., 2014, Lu et al., 1993, Mihalick et al., 2000, Passe and Walker, 1985, Petrulis and Eichenbaum, 2003, Rinberg et al., 2006, Slotnick et al., 1991, Uchida and Mainen, 2003). Automatic behavioral systems have been developed for studying innate olfactory behaviors (Qiu et al., 2014). Olfactory behavioral testing has been developed in head-fixed rodents and greatly facilitates the understanding of neural circuits underlying olfaction (Verhagen et al., 2007; Wesson et al., 2008; Shusterman et al., 2011; Kato et al., 2013; Boyd et al., 2015) and odor-based cognition (Komiyama et al., 2010; Liu et al., 2014; Gadziola et al., 2015). However, fully automatic training systems for odor-based cognitive behaviors were not available for head-fixed mice.

We therefore designed a high-throughput automatic training system (HATS) for olfactory behaviors in head-fixed mice. Using the automatic step-by-step training procedures, we trained mice to perform olfactory delayed non-match to sample (DNMS), delayed paired association (DPA), Go/No-go (GNG), and GNG reversal tasks. Mice reached stable levels of performance within several days in the tasks. HATS can be an important tool in our understanding of the neural-circuitry mechanisms underlying odor-based cognitive behaviors.

## Material and Methods

### Animals

Male adult C57BL/6 mice (SLAC, as wild-type) were used for the current study (8-40 weeks of age, weighted between 20 to 30 g). Wild-type mice were provided by the Shanghai Laboratory Animal Center (SLAC), CAS, Shanghai, China. Mice were group-housed (4-6/cage) under a 12-h light-dark cycle (light on from 5 a.m. to 5 p.m.). Before behavioral training, mice were housed in stable conditions with food and water ad libitum. After the start of behavioral training, the water supply was restricted. Mice could drink water only during and immediately after training. Care was taken to keep mice body weight above 80% of a normal level. The behavioral results reported here were collected from a total of 25 wild-type mice. All animal studies and experimental procedures were approved by the Animal Care and Use Committee of the Institute of Neuroscience, Chinese Academy of Sciences, Shanghai, China.

### Animal surgery

Mice were anesthetized with analgesics (Sodium pentobarbital, 10mg/mL, 80 mg/kg body weight) before surgery. All surgery tools, materials, and experimenter-coats were sterilized by autoclaving. Surgery area and materials that cannot undergo autoclaving were sterilized by ultraviolet radiation for more than 20 minutes. Aseptic procedures were applied during surgery. Anesthetized mice were kept on a heat mat to maintain normal body temperature. Scalp, periosteum, and other associated soft tissue over skull were removed. Skull was cleaned by filtered artificial cerebrospinal fluid (ACSF) with cotton applicators. After skull was dried out, a layer of tissue adhesive was applied on the surface of the skull. A steel plate was placed on the skull and then fixed by dental Cement.

### Behavior setups

HATS was composed of a mouse containing, head-fix, odor delivery and reward delivery, Arduino based control, and data acquisition units (diagram in Figure 1A, photo in Figure 1B). All valves and motors were controlled by Arduino based processors and customized software. The 3d printing files, a step by step instruction for hardware assembling, the source code for behavior training and the data acquisition source code were publically available (https://github.com/wwweagle/serialj; https://github.com/jerryhanson/frontiers/)

**FIGURE 1.**
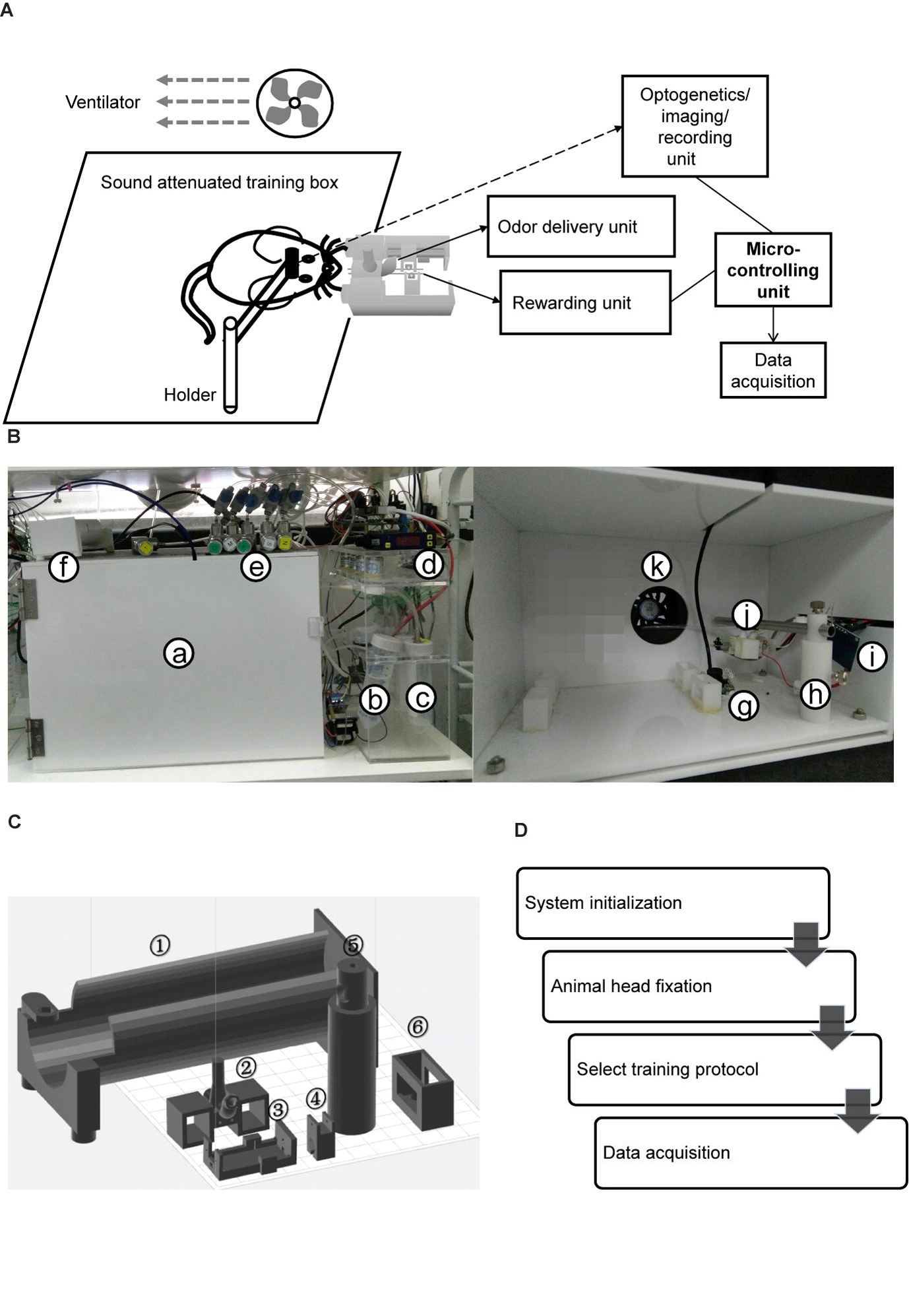
Components and operational processes for HATS. **(A)** Schematics showing the components of HATS. **(B)** Photos of HATS hardware. a, sound-attenuated box; b-c, odor containers; d, flow meter; e, needle valve; f, training tube for restraining mouse body; g, camera; h, holder for the odor- and water-delivery unit and motors; i, capacitance detector for licking; j, 3D-printed odor and water delivery unit; k, ventilator. **(C)** 3D-printed components. 1, mouse-body tube; 2, a socket for odor-delivery tubes; 3-4, slots for the motor of the moveable water port; 5, a holder for the odor- and water-delivery unit and motors; 6, a socket for the training tubes. **(D)** Schematics of the operational processes for HATS.

Three-dimensional printing technique was used to generate the small components in the system (Figure 1C). The training tube was used to maintain the relative position of mouse body to the water- and odor-delivery ports. The motor slot held a direct-current motor to move the water port forward or backward. The water tube slot held a metal needle with a blunt tip, from which mice obtained water as a reward. The odor-tube slot connected the odor tube from the odor-delivery unit.

A movable water port was connected to a peristaltic pump, which was controlled by an Arduino board. The volume of water reward was controlled by changing the duration of the output signal to peristaltic pump from the Arduino board. Peristaltic pumps of different setups were calibrated for the stable volume of water delivery in each trial (5 ± 0.5 μL).

Water- and odor-delivery units were both controlled by an Arduino board. During behavior training, detailed timing information of events was sent back to the computer via the USB-simulated serial-port interface and stored by a customized Java program. The stored events included an odorant valve on/off, peristaltic pump on/off, and licking start/end. Licking event was detected by a capacity detector. Infrared LED-based licking detectors were used for electrophysiological recording if required. An infrared camera was placed under the water port to monitor behavioral states of mice.

### Olfactometer

The olfactometer was designed to efficiently and reliably mix and deliver odor. Air source was a pump that provided air flow with the flow rate of ~120 L/min. The filter was applied to eliminate moisture and dust. Eight training setups shared one set of pump and filter. For each setup, pure air with the flow rate of 2 L/min is constantly delivered to mice during the entire process. The air input to each air route could be turned on and off by a manual valve (labeled as “M” in Figure 2A and 2B). The flow rate was adjusted by a needle valve (labeled as “V” in Figure 2A and 2B). As shown in the Figure 2B, one type of odorant in liquid state was stored in one airtight bottle. The air-in tube was placed right above the surface of the liquid odorant. Two-way solenoid valves were used to switch the odor to either mouse or flow mater. In the standby state (no odor was delivered, Figure 2A), the valve to odorant bottle (labeled as “O”) was closed, and that to the flow meter (labeled as “F”) was opened. Therefore, no odor will be mixed with pure air and delivered to the mouse. In the working state that odor was delivered (Figure 2B), “O” was open and “F” was closed. Therefore odor was mixed with constant air and delivered to the mouse. Four kinds of odorants were used in the behavior tasks, 1-Butanol, Methyl butyrate, Hexanoic acid, and Octane. The relative volume ratios of these odorants in the pure air were 10%, 2.5%, 15% and 5%, respectively. The difference was due to the distinct evaporation pressure of different odorant molecules at room temperature (see Table 1 for detailed rising/decay and residual time of the odorants). The odor tubes after “O” valves and before mixture chamber had an inner diameter of 0.5 mm. The odor tube for constant air before mixture chamber had an inner diameter of 2.5 mm.

**FIGURE 2.**
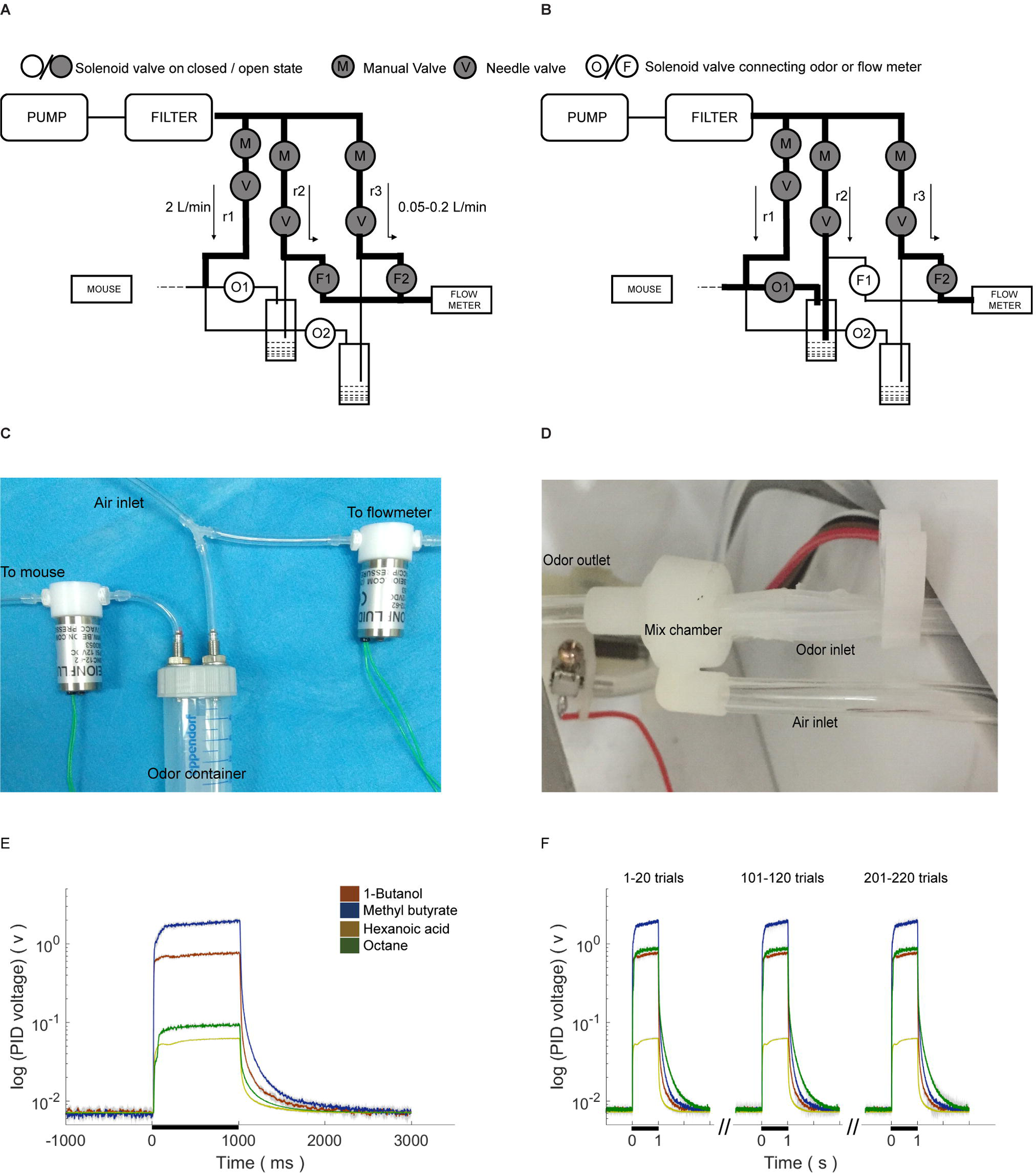
Design, implementation, and reaction time of the olfactometer. **(A)** Schematic showing the “standby” condition of the olfactometer. Diagram for the two-odor delivery unit was shown. The flow meter was designed for monitoring potential system failure. The flow rate was labeled as the numbers with the unit “L/min.” Arrows indicated for the direction of air flow. **(B)** Schematic showing the “working” condition when one odor was delivered (through “r2”). Reduction of the readout from the flow meter indicated for normal operation. **(C)** Photo of the flow-controlling unit for the olfactometer. **(D)** Photo of the tubing unit and mixing chamber. Thin tubes were used for fast reaction for odor delivery. Mixing chamber was designed for a maximal mixture of pure air (from “r1” in **B**) and the delivered odor (from “r2” in **B**). **(E)** Fast response of the olfactometer. Readout from PID was plotted in the log scale for main figure and linear scale for inset (Mean ± SEM, standard error of the 0mean, unless stated otherwise; calculated from odor application of 200 trials). Rising/decay time constant and time with residual-odor were shown in Table 1. **(F)** Odor stability across trials.

**TABLE 1.** Rising and decay properties of odorants. Time was in a millisecond.

| Odorant Name | Relative Volume Ratio in Air (%) | Rising Latency (95% of peak) | Decay Constant Time (1/e of peak) |
| --- | --- | --- | --- |
| 1-Butanol | 10 | 18 ± 1 | 20 ± 1 |
| Methyl butyrate | 2.5 | 17 ± 1 | 22 ± 1 |
| Hexanoic acid | 15 | 31 ± 1 | 41 ± 1 |
| Octane | 5 | 71 ± 1 | 31 ± 1 |

(mean ± standard error of the mean)

### Behavior training

#### Water restriction

Mice were allowed at least seven days for recovery after surgery for head-plate implantation. Before the start of formal training, mice were water restricted for 48~72 hours, in which licking for water was allowed (less than 1.0 ML per day, exact amount was not monitored). Throughout the training, the daily intake of water was at least 0.6 ML per day (as in Guo et al., 2014) and typically 1.0 ML per day. Body weight was closely monitored and a steady increase in body was observed after initial decrease following 24 hour restriction.

#### Habituation phases

The habituation phase started thirty minutes before the start of the training phase and only occurred once. A training tube was placed into the home cage. Mice could explore the tube freely to be familiar with it. This step was designed to decrease the stress level of mice on the first day.

#### Automatic licking teaching phase

This phase was designed to teach mice to lick freely from the water tube. A mouse was fixated on the head plate to a holding bar connected to the training tube. The animals were transferred from homecages to the apparatus and headfixed manually by experimenters. The total time spent in transition was less than a minute. Then the training tube was placed into and fixated to sliding sockets in the sound-attenuated box (the typical decrease from background noise was 15dB). Initially, the tip of the water port was placed five millimeters away from the mouse mouth. By using a program-controlled movable water port, the initiation of a teaching bout was associated with the forward movement of the water port. During each day, this phase was divided into three bouts to facilitate the association between movement of the water port and delivery of water. In each bout, water port moved forward firstly to seduce mouse to lick. After two seconds, water port will be reset back to the original place. Once mouse licked, one water drop (volume of ~5 μL) was delivered for every three licks. This bout ended when mice did not lick continuously for two seconds, or rewarded size is larger than 200 μL from this bout. The daily reward size could vary between each mouse (typically 0.6 ML and less than 1.0 ML). This phase lasted for three days. Mice stayed in training apparatus for 1-2 hours per day in all training phases.

#### Automatic shaping phase

This phase was designed to teach mice to lick for water only in the response window, which was from 0.5 to 1.5 sec after the offset of the second odor delivery. Only rewarded condition was applied, which were non-matched pairs for DNMS task, paired odors for DPA task, or go cue for GNG task, respectively. Mice could lick in response window to trigger water reward from every trial. During this phase, water port may or may not move while water was delivered. If mice missed several trials, lick-teaching would resume, in which the water port was moved forward and water was delivered during the response window. The reward in lick-teaching was program-controlled and was not triggered by lick. Two types of trials were defined for this phase, the self-learning (Figure 5, left) and program-teaching (Figure 5, right) trials, which switched automatically under the condition introduced below. The water port was moved forward during the response window in the program-teaching trials, in which the water delivery was automatic without triggered by licks. In the self-learning trials, however, reward delivery was licking-triggered, and water port did not move. The condition for switching from self-learning to program-teaching trials was that mice missed five times within 30 trials or missed during the last program-teaching trial. The condition for switching from program-teaching trial to self-learning trial is that mice licked in response window and obtained a reward from the last teaching trial. Daily shaping phase ended when mice performed 100 hit trials in total. This phase lasted for three days.

### Full task training phase

#### DNMS task training

In the DNMS task, a sample odor was delivered at the start of a trial, followed by a delay period (4-5 seconds) and then a test odor, same to (matched) or different from (non-matched) the sample (Figure 6). Two kinds of odorants were used in DNMS task, 1-Butanol, and Methyl butyrate. The relative volume ratios in the pure air were 10% and 2.5%, respectively. Odor-delivery duration was one second. Mice were trained to lick in the response window in non-match trials. The response window was from 0.5 to 1.5 sec after the offset of the second odor delivery. Licking events detected in the response window in the non-match trials were regarded as Hit and will trigger instantaneous water delivery (a water drop around 5 μL). The false choice was defined as detection of licking events in the response window in the match trials. Mice were not punished in the False Choice trials. Mice were neither punished nor rewarded for the Miss (no-lick in a non-match trial) or the Correct rejection (CR, no-lick in a matching trial) trials. Behavioral results were binned in blocks of 24 trials. There was a fixed inter-trial interval of 10 seconds between trials. After training ended each day, mice were supplied with water of at least 300 μL and up to 1 mL daily intake. This phase lasts for four to five days. The well-trained criterion was set to the existence of three continuous correct-rates larger than 80%, calculated using a sliding window of 24 trials. The reason to use 24 trials as a block is to maintain the consistency of different trial types between different tasks, with the need to be commonly divided by four and eight types of odor sequence for different tasks (4 for DNMS, 4 for DPA). It was intended to facilitate the compairsion of the performance in the different tasks in the the current study. It can be easily modified according to the needs of.

#### DPA task training

For the DPA task, a sample and a test odor were delivered, separated by a delay period (Figure 7). Four kinds of odorants were used, 1-Butanol (S1), Methyl butyrate (S2), Hexanoic acid (T1) and Octane (T2). The relative volume ratios in pure air were 10%, 2.5%, 15% and 5%, respectively. Odor delivery duration was one second. Delay period between two odors in a trial was 8-9 seconds. Response window was set to 0.5-1 second after the offset of the test odor in a trial. Mice were trained to lick to obtain water reward only after the paired trials (S1-T1 or S2-T2). Licking events detected in the response window in paired trials were regarded as Hit and will trigger instantaneous water delivery. The false choice was defined as detection of licking events in the response window in non-paired trials (S1-T2 or S2-T1), and mice were not punished in False Choice trials. Mice were neither punished nor rewarded for Miss (no-lick in the paired trial) or Correct rejection (CR, no-lick in a non-paired trial) trials. Behavioral results were binned in blocks of 24 trials. There was a fixed inter-trial interval of 16 seconds between trials. After training ended each day, mice were supplied with water of at least 300 μL and up to 1 mL daily intake. This phase lasts for four to five days. The well-trained criterion was set to the existence of three continuous correct-rates larger than 80%, calculated using a sliding window of 24 trials.

#### GNG and GNG reversal task training

For the GNG task, mice were trained to lick for water only after the Go cue but not No-go cue. Hexanoic acid and Octane were used as Go and No-go cues, respectively. The relative volume ratios in the pure air were 15% and 5%, respectively. Odor-delivery duration was one second. Response window was 0.5-1.5 second after the offset of a cue. Licking events detected in the response window in Go trials were regarded as Hit and triggered instantaneous water delivery. The false choice was defined as the detection of licking events in the response window in No-go trials. Mice were not punished in the False Choice trials. Mice were neither punished nor rewarded for the Miss (no-lick in a Go trial) or the Correct rejection (CR, no-lick in a No-go trial) trials. Behavioral results were binned in blocks of 24 trials. There was a fixed inter-trial interval of 5 seconds between trials. After training ended each day, mice were supplied with water of at least 300 μL and up to 1 mL daily intake. This phase lasts for three days. The well-trained criterion was set to the existence of three continuous correct-rates larger than 80%, calculated using a sliding window of 24 trials.

In the third day of training, the GNG reversal task began, in which the odor-reward relationship was reversed.

#### Data analysis

The performance of the correct rate (referred to as “performance” in labels of figures) of each bin was defined by:

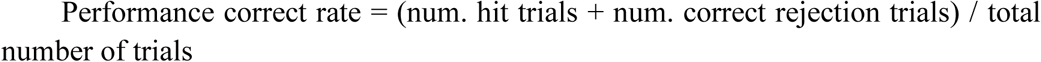

Hit, False choice, and Correct rejection rates were defined as follows:

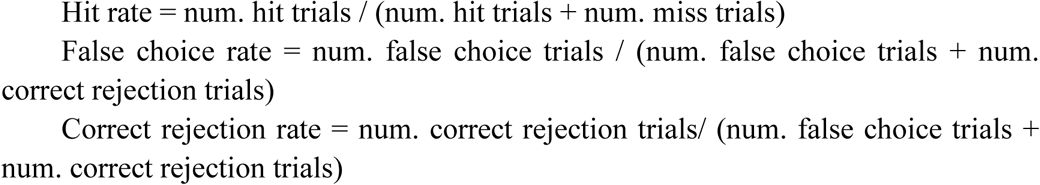

Mean correct rate (CR rate/ FA rate) was calculated as an averaged correct rate (CR rate / FA rate) between different mice.

Error bars from the mean value of the correct rate (CR rate / FA rate) was calculated by the standard error of the mean. N represents the number of mice.

The licking rate was calculated as lick numbers within each time bin (bin size:100 ms). The curve was smoothed by smooth function from Matlab with a span size of 5 bin.

Discriminability (*d’*) was defined by:

*d’ =* norminv (Hit rate)-norminv (False choice rate). The norminv function was the inverse of the cumulative normal function. Conversion of Hit or False choice rate was applied to avoid plus or minus infinity (Macmillan and Creelman, 2005). In conversion, if Hit or False choice rate was equal to 100%, it was set to [1-1/(2n)]. Here, n equals to a number of all possible Hit or False choice trials. If Hit or False choice rate was zero, it was set to 1/(2n).

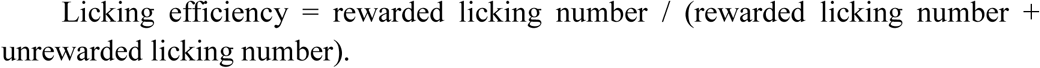

A number of trials to criterion was calculated as the trial numbers before reaching 80% correct rate for 24 consecutive trials. “NRC” in Figure 6-8 represented Not Reaching Criterion, which indicated that mice did not reach the above criterion for that day.

## Results

### Overview of Hardware, Software, and protocol

In our previous study (Liu et al., 2014), mice were manually taught to lick for water and shaped for a DNMS task. The goal of the current study was to allow fully automatic training. The only things human operators need to perform were to fixate mice onto head-fix bars, close doors of training boxes, and run computer software controlling training protocols. The current study fulfilled the goal by designing HATS for olfactory and odor-based cognitive behavior in head-fixed mice. HATS was composed of a mouse containing, head-fix, odor- and water-delivery, Arduino based control, and data acquisition units (diagram in Figure 1A, photo in Figure 1B, 3D-printed parts in Figure 1C). Optogenetic, chemogenetic, recording, and imaging methods can be easily integrated into HATS. All valves and motors were controlled by Arduino based processors and customized software. The daily routine was composed of system adjustment, head-fixation of mice, choosing a protocol, and training mice a given behavior (Figure 1D).

### Fast odor delivery

In studying olfactory behaviors, it is critical to have fast rise and decay for odor delivery.

Our olfactometer exhibited fast response and stable performance. The reaction time constant for the onset of these odors was between 11-71 ms (**Table 1**), measured with a photoionization detector (PID). Another key parameter was the time constant for decay after the offset of the odor-delivery unit, which was especially important in working memory-related tasks. The current odor-delivery unit exhibited fast decay (time constant: 20-41 ms, Figure 2C, Table 1). Moreover, odor concentration remained stable following more than 200 trials of odor delivery (Figure 2F), which was important for behavioral and recording experiments.

### Automatic training protocol

To achieve fully automatic training, we developed a step-by-step training protocol. The protocol was separated into two preparatory steps (water deprivation and habituation) and three training phases (lick-teaching, shaping, and learning, Figure 3A).

**FIGURE 3.**
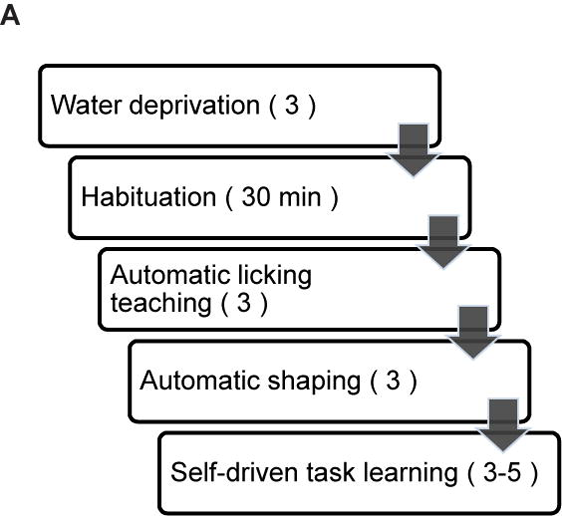
Step-by-step automatic training procedure. **(A)** Step-by-step automatic training procedure. Duration in a given step was labeled to the right.

The first step of training was to automatically teach licking freely from water tube (Figure 4A-B). Moveable water port (Figure 4C-E) was located 5 mm away from the mouth of a mouse. The flow chart of the lick-teaching protocol was plotted in Figure 4A. At the start of a teaching bout, water port would deliver 10 μL water and then moved forward until contacting the mouth, thus encouraging the licking. If mouse licked, 4 μL water would be rewarded for every three licks. After no licking was detected for consecutive 2 sec or water of 200 μL was delivered, one bout of teaching was completed, and the water port was moved back to the initial position. The teaching bout was repeated for several times until water of 400 μL was rewarded in total. The volume of water rewarded in each day was plotted in Figure 4B.

**FIGURE 4.**
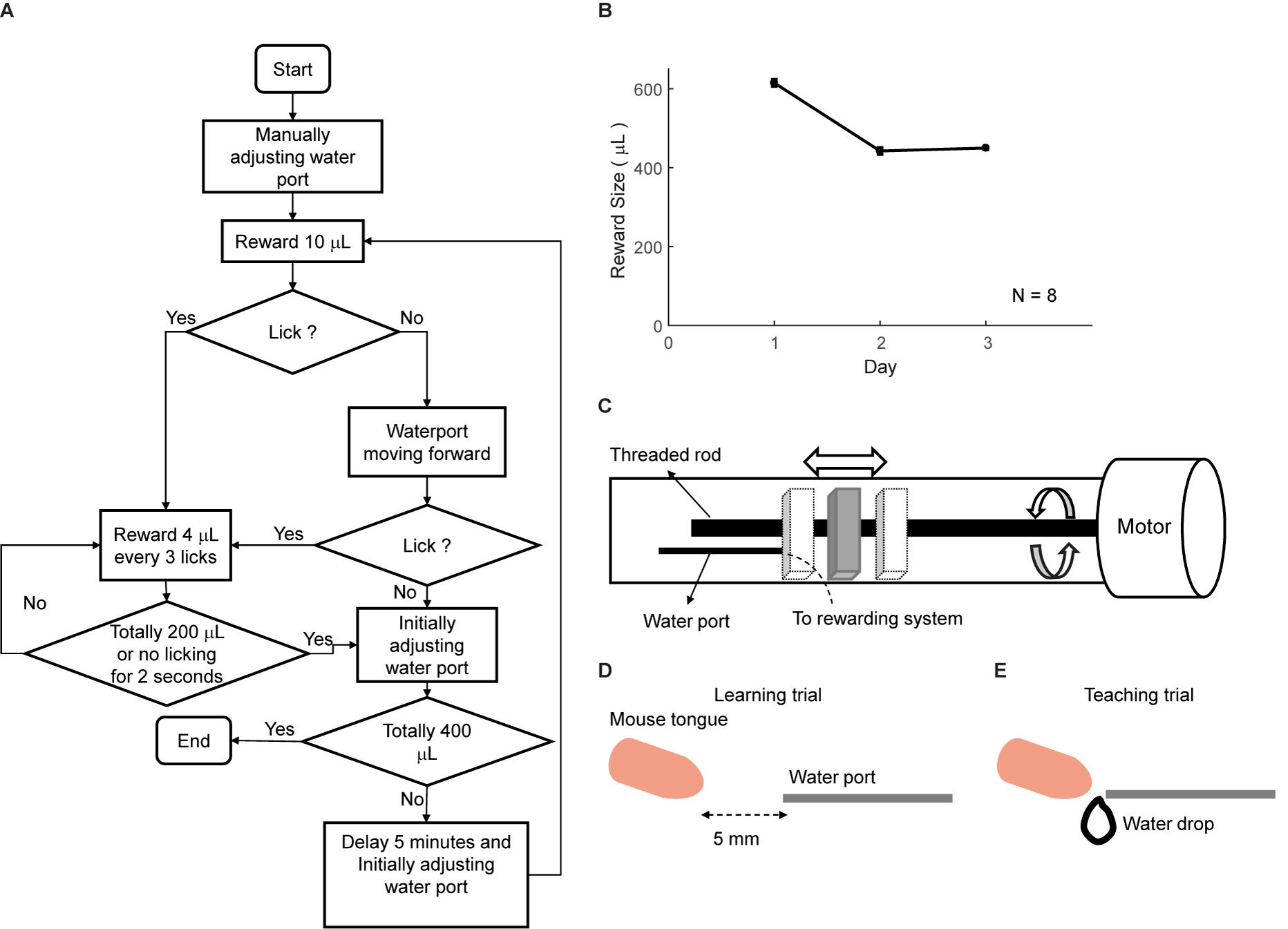
Automatic lick-teaching protocol. **(A)** Flow chart for the automatic lick-teaching protocol. **(B)** Daily consumed water volume in the lick-teaching phase. **(C)** Diagram of the moveable water port. **(D-E)** Diagram showing relative position between water port and mouse mouth in self-learning (**D**) and teaching (**E**) phases.

The second step of training was shaping for a specific task. This phase was designed to allow mice to be familiar with the temporal structure of the tasks and the involved sensory stimuli, without experiencing the full task. In shaping, only the trials with water reward were applied. Specifically, for DNMS task, only non-matched odor pairs were applied to mice (Figure 5A-B). For DPA task, only paired trials were applied. For GNG task, only Go cue was applied. Two types of trials were designed, self-learning and teaching trials. In self-learning trials, water delivery was triggered by licking in the response window (Figure 5C left box). In teaching trials, water port moved forward and delivered water automatically during response window (Figure 5C right box). These two types of trials were designed to switch automatically. The condition for switching from learning to teaching trial was that mice missed five trials in 35 trials. The condition for switching from teaching to learning trial was that mice licked within the response window in the last teaching trial. Daily shaping phase ended when mice performed 100 hit trials in total. This phase lasted for three days.

**FIGURE 5.**
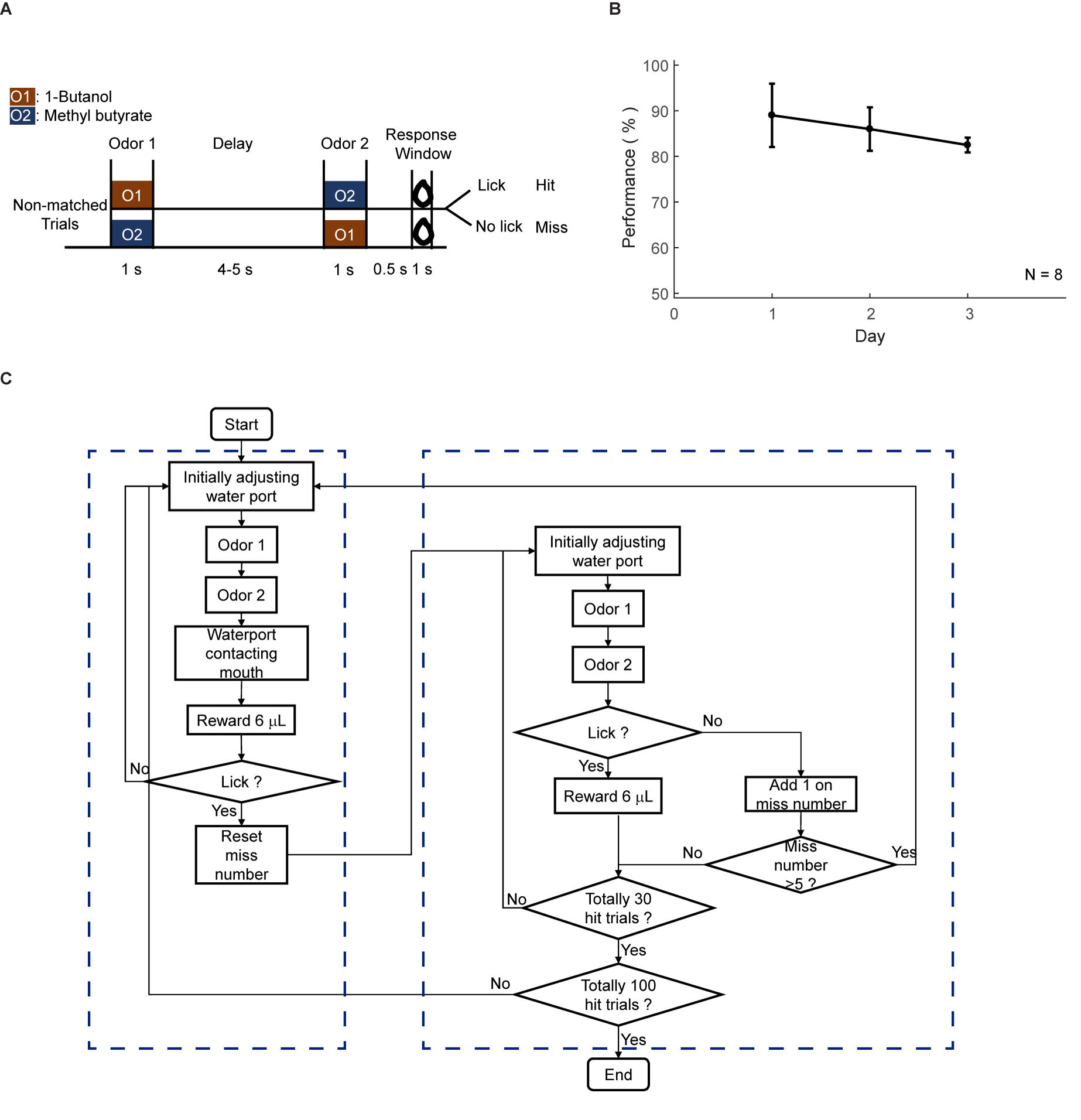
Automatic shaping protocol. **(A)** Design paradigm and time line for the DNMS shaping. Only non-matched trials were applied. **(B)** Licking performance in the shaping phase. **(C)** Flow chart for the DNMS shaping. Left: self-learning trials. Right: teaching trials.

### Training the DNMS task

We trained eight head-fixed mice to perform an olfactory DNMS task (Liu et al., 2014) (Figure 6A). In this design mouse needed to temporally maintain information during the delay period before behavioral choices and motor planning. After the shaping protocol, we added the non-rewarded matched trials, which induced false choice and reduced performance to chance level (Figure 6B). Gradually the performance, correct rejection, and discriminability (*d’*) progressively increased, whereas the hit rate remained at a ceiling level (Figure 6B-E). After the training of five days (600 trials), the performance showed significant increase (ANOVA, p<0.0001, F=775.89). Mice experienced a certain level of relearning each day, with a decreased number to criteria (defined as a correct rate above 80% in 24 consecutive trials) each day through learning (Figure 6F). Most of the licking responses were associated with non-match odor and expectation of water reward (Figure 6G). There were licks associated with the first odor delivery in the early phase of learning (Figure 6G black curve), which were declined through learning (Figure 6G blue curve). Also, the licking efficiency (defined as the ratio of successful licks resulting water reward) was increased progressively through learning (Figure 6H).

**FIGURE 6.**
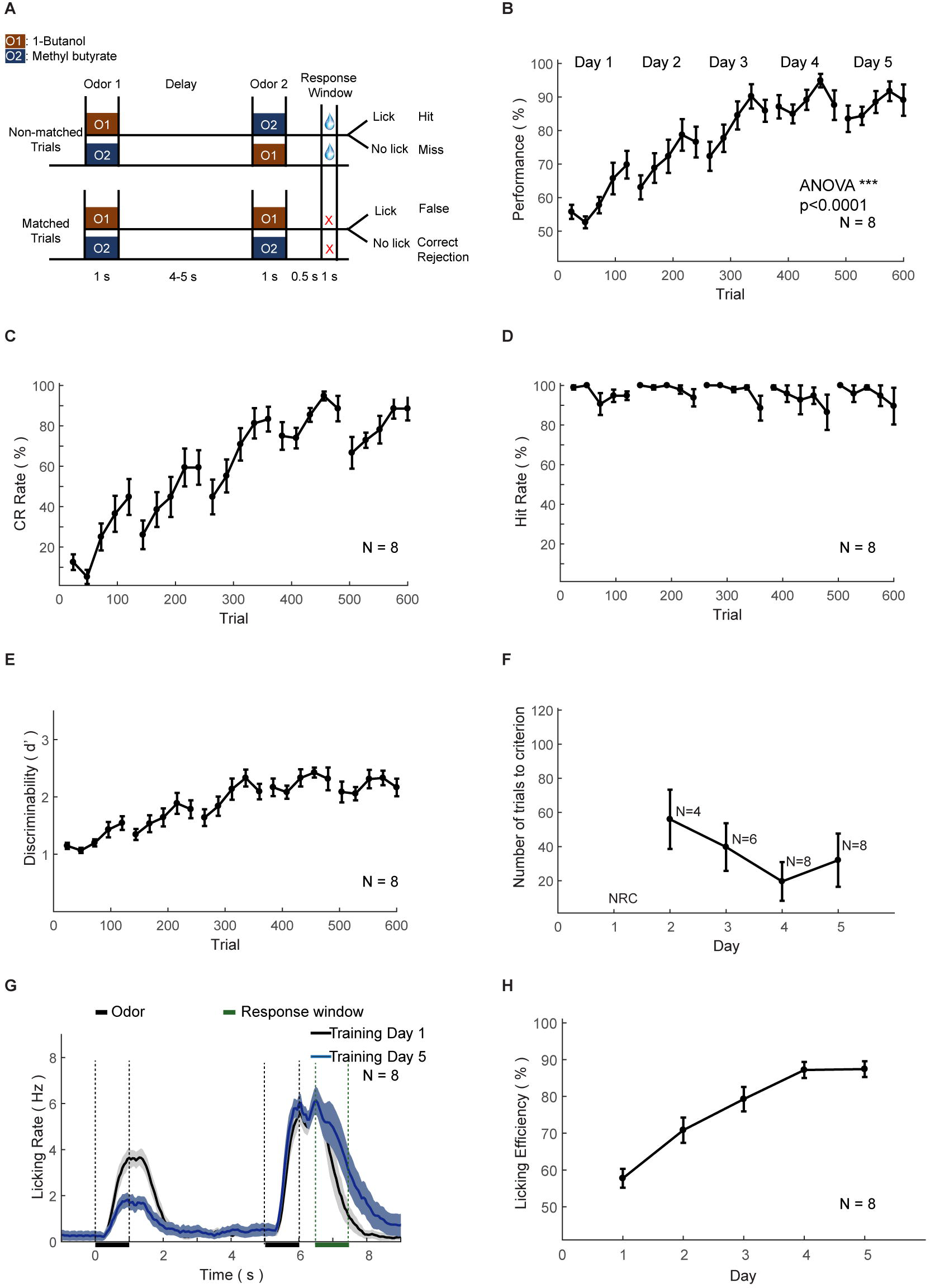
Automatic DNMS training protocol and behavioral results. **(A)** Design paradigm and time line for the DNMS training. Both non-matched and matched trials were applied. **(B)** Performance of mice in the DNMS training phase. Bin size: 24 trials. **(C-E)** Correct rejection (CR) rate, hit rate, and *d’* in the DNMS training, respectively. **(F)** Re-learning in each day of the DNMS training, measured by the number of trials to criterion (defined as more than 80% performance in 24 consecutive trials). NRC, not reaching criteria. Mice that were NRC in the 2^nd^ and 3^rd^ days were not included. **(G)** Licking rates for training day 1 and 5. **(H)** Licking efficiency in the DNMS training. Licking efficiency was defined as the ratio of successful licks resulting water reward.

### Training the DPA task

The second set of head-fixed mice was trained to perform an olfactory DPA task (Figure 7A). As in the DNMS task, the performance, correct rejection, and discriminability (*d’*) progressively increased, whereas the hit rate remained at ceiling level (Figure 7B-E). After the training of five days (600 trials), the performance showed significant increase (ANOVA, p<0.0001, F=1139.03). Mice also experienced a certain level of relearning each day (Figure 7F). Most of the licking responses were associated with paired odor and expectation of water reward (Figure 7G). There were licks associated with the first odor delivery in the early phase of learning (Figure 7G black curve), which were declined through learning (Figure 7G blue curve). Such early licks associated with the sample odor were lower than that in the DNMS task. The licking efficiency also increased progressively through learning (Figure 7H).

**FIGURE 7.**
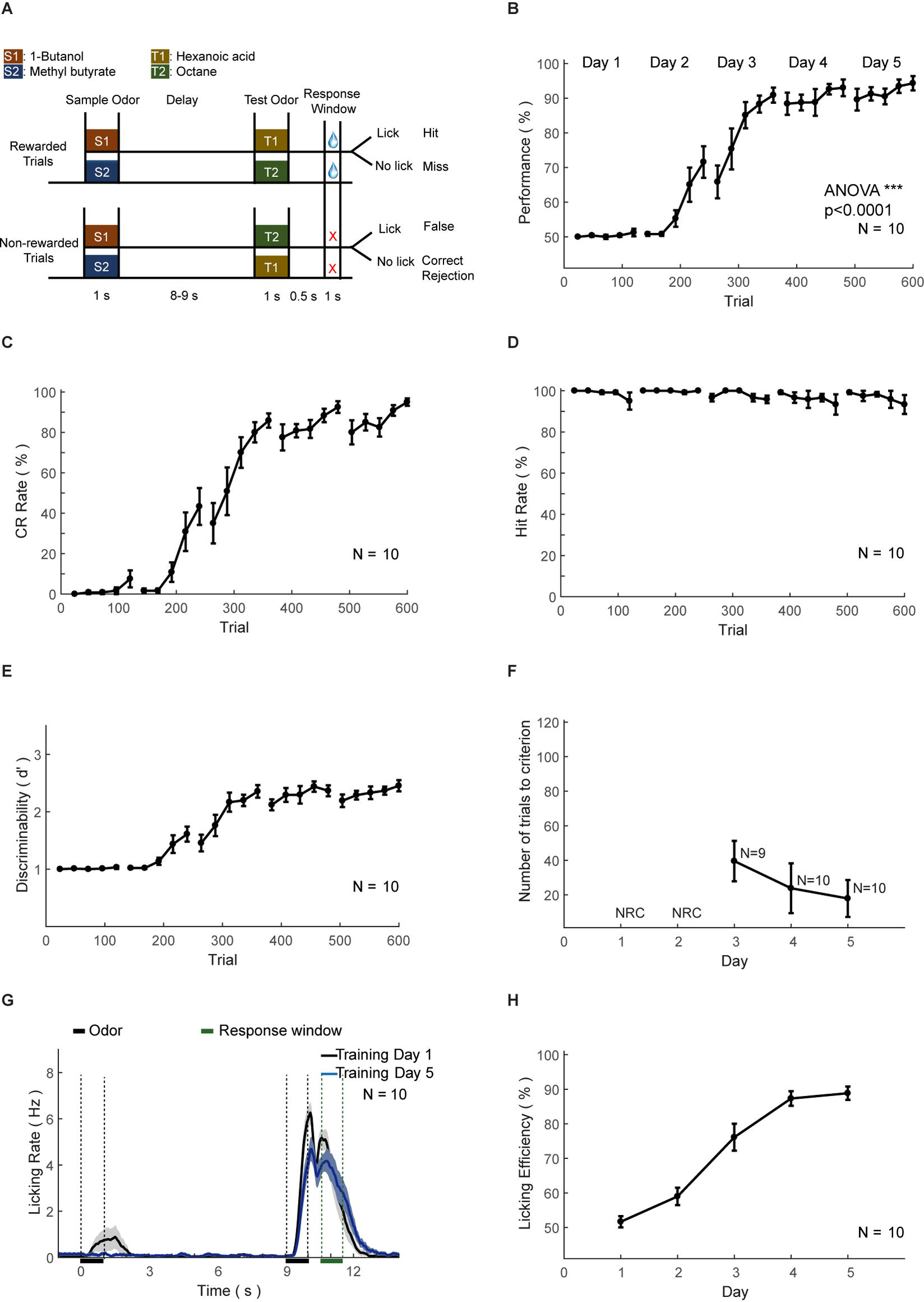
Automatic DPA training protocol and behavioral results. **(A-H)** As in Figure 6 A-H.

### Training the GNG and reversal tasks

The third set of head-fixed mice was initially trained to perform an olfactory GNG task (Figure 8A above), then subsequently sensory-cue reversal task (Figure 8A below). The performance, correct rejection, and discriminability (*d’*) progressively increased, whereas the hit rate remained at ceiling level (Figure 8B-E). After the training of two days (200 trials), the performance showed significant increase (ANOVA, p<0.0001, F=3455.17). Mice also experienced a certain level of relearning each day (Figure 8F). Most of the licking responses were associated with paired odor-pair and expectation of water reward (Figure 8G). The licking efficiency also increased progressively through learning (Figure 8H). After two-days of GNG training, the odor-reward relationship was reversed (Figure 8A below). The performance, correct rejection, discriminability (*d’*), and licking efficiency were decreased initially, and then progressively increased (Figure 8B-E). The hit rate remained at ceiling level (Figure 8D) and relearning was evident from the number of trials to criteria (Figure 8F).

**FIGURE 8.**
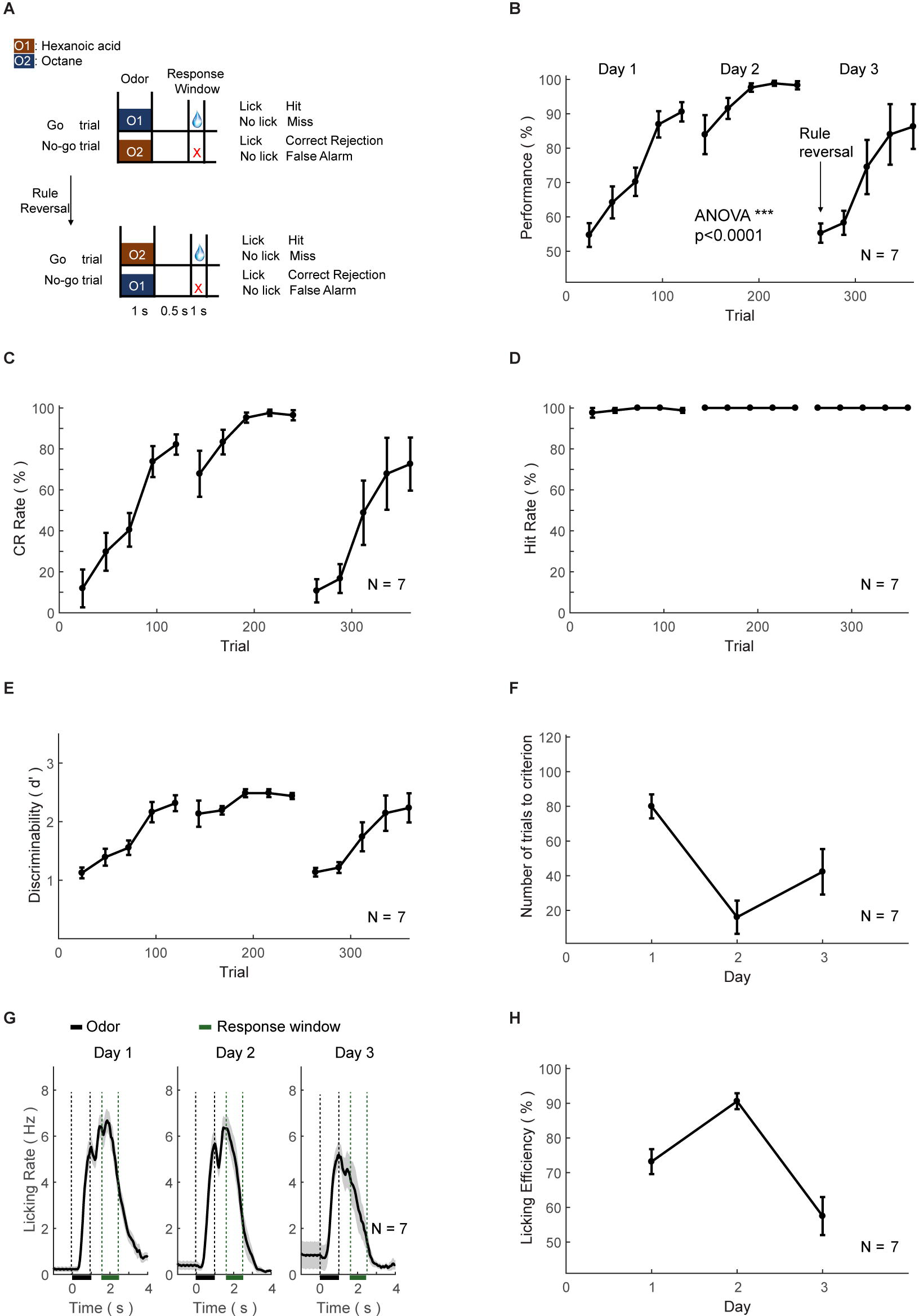
Automatic GNG and reversal training protocol and behavioral results. **(A-H)** As in Figure 6 A-H.

## Discussion

Automated, quantitative, and accurate assessment of behaviors is critical for understanding mechanisms underlying cognition. Here we presented HATS, a new integrated hardware and software system that combined fast olfactometer, 3D-printed components, step-by-step automatic training, for automatic training of cognitive behaviors in head-fixed mice. The robustness of the system was validated in multiple olfactory and odor-based tasks. The involved tasks require cognitive abilities including working memory (Fuster, 1997; Baddeley, 2012), decision making (Gold and Shadlen, 2007; Lee et al., 2012), and reversal of learnt rules (Bunge and Wallis, 2008), all of which are required in more naturalistic environment and vital for survival.

An obvious limitation is that free-moving mice cannot be trained with HATS. Another limitation is that HATS only monitor the lick as behavioral readouts, therefore is more suited for large-scale screening of optogenetic. Although the head-movement was restrained in the current design, one would like to monitor the muscles controlling head or chewing movement to further eliminate the potential artifacts in electrophysiological recording. To obtain deep understanding of neural circuit underlying these behavior, one would also like to integrate more monitoring systems for behavioral events, such as sniffing (Kepecs et al., 2007; Verhagen et al., 2007; Wesson et al., 2008; Shusterman et al., 2011; Deschenes et al., 2012; McAfee et al., 2016), pupil size (Reimer et al., 2014; McGinley et al., 2015; Vinck et al., 2015; Bushnell et al., 2016; Reimer et al., 2016), and whisker movement (Orbach et al., 1985; Friedman et al., 2006; Birdwell et al., 2007; O’Connor et al., 2010; Deschenes et al., 2012; Petreanu et al., 2012; Moore et al., 2013).

In designing HATS, we tried to fasten the training history, therefore aiding the dissection of neural circuit. However, this fast training in animals would only sufficiently model fast learning in humans. Indeed, many human behaviors and human learning are slow in learning and require extensive training, such as fine motor skill (*i.g.*, driving, playing piano) and sensory discrimination (*i.g.*, wine tasting). Thus, automations achieved in HATS have limitations to what kinds of behavioral and neural processes are being effectively modeled.

Nevertheless, HATS allowed for rapid, automated training of cognitive behaviors across diverse experimental designs. Our approach can also support high-throughput behavioral screening. In summary, the newly developed HATS is well-suited for circuitry understanding of odor-based cognitive behavior.

## Acknowledgments

The work was supported by the Strategic Priority Research Program of the Chinese Academy of Sciences (Grant No. XDB02020006), the Chinese 973 Program (2011CBA00406), the Instrument Developing Project of the Chinese Academy of Sciences (Grant No. YZ201540), the Key Research Project of Frontier Science of the Chinese Academy of Sciences (Grant No. QYZDB-SSW-SMC009), China – Netherlands CAS-NWO Programme: Joint Research Projects, The Future of Brain and Cognition (153D31KYSB20160106), the Key Project of Shanghai Science and Technology Commission (No.15JC1400102, 16JC1400101), the National Science Foundation for Distinguished Young Scholars of China (31525010, to C.T.L.), the General Program of Chinese National Science Foundation (31471049), the State Key Laboratory of Neuroscience, and CAS Hundreds of Talents Program (2010OHTP04, to C.T.L.).

